# Targeted stimulation of human orbitofrontal networks disrupts outcome-guided behavior

**DOI:** 10.1101/740399

**Authors:** James D. Howard, Rachel Reynolds, Devyn E. Smith, Joel L. Voss, Geoffrey Schoenbaum, Thorsten Kahnt

**Affiliations:** Department of Neurology, Northwestern University, Feinberg School of Medicine, Chicago, IL 60611, USA; Department of Medical Social Sciences, Northwestern University, Feinberg School of Medicine, Chicago, IL 60611, USA; Department of Psychiatry and Behavioral Sciences, Northwestern University, Feinberg School of Medicine, Chicago, IL 60611, USA; National Institutes on Drug Abuse, Intramural Research Program, Baltimore, MD 21224, USA; Department of Psychology, Northwestern University, Weinberg College of Arts and Sciences, Evanston, IL 60208, USA

**Keywords:** Orbitofrontal cortex, reward, decision-making, inference, devaluation, transcranial magnetic stimulation, outcome-guided behavior, functional connectivity

## Abstract

Outcome-guided behavior requires knowledge about the current value of expected outcomes. Such behavior can be isolated in the reinforcer devaluation task, which assesses the ability to infer the current value of rewards after devaluation. Animal lesion studies demonstrate that orbitofrontal cortex (OFC) is necessary for normal behavior in this task, but a causal role for human OFC in outcome-guided behavior has not been established. Here we used sham-controlled non-invasive continuous theta-burst stimulation (cTBS) to temporarily disrupt human OFC network activity prior to devaluation of food odor rewards in a between-subjects design. Subjects in the sham group appropriately avoided Pavlovian cues associated with devalued food odors. However, subjects in the stimulation group persistently chose those cues, even though devaluation of food odors themselves was unaffected by cTBS. This behavioral impairment was mirrored in changes in resting-stated functional magnetic resonance imaging (rs-fMRI) activity, such that subjects in the stimulation group exhibited reduced global OFC network connectivity after cTBS, and the magnitude of this reduction was correlated with choices after devaluation. These findings demonstrate the feasibility of indirectly targeting the human OFC with non-invasive cTBS, and indicate that OFC is specifically required for inferring the value of expected outcomes.

## INTRODUCTION

To make adaptive choices, organisms must anticipate the value of expected outcomes. In the face of continually changing motivational states and external contingencies, this requires the ability to infer the current value of specific outcomes on-the-fly, without the need for new learning [1, 2]. For example, when perusing the menu at a new restaurant, we can readily infer how much we will like each option, and make a choice without having to try each one first. Such inference, or mental simulation, is a hallmark of outcome-guided behavior, distinguishing it from behavior that can be based on first-hand experience [3, 4].

Decisions that require inference can be isolated in the reinforcer devaluation paradigm, in which responses to a predictive cue are probed after selective devaluation of an associated outcome [5]. Experiments in rodents and non-human primates demonstrate that inactivation of the orbitofrontal cortex (OFC) results in continued responding to Pavlovian cues predicting a devalued outcome, indicating an inability to infer its new value [6–13]. Yet, while neuroimaging studies show a correlation between human OFC activity and updated reward expectations in devaluation tasks [14–16], definitive evidence in support of a causal role for human OFC in outcome-guided behavior is lacking.

Activity in the human brain can be modulated non-invasively using transcranial magnetic stimulation (TMS)[17]. Yet, due to its anatomical location, the OFC is not directly accessible to surface stimulation techniques such as TMS, making it difficult to test the causal role of OFC in inference-based decisions in healthy humans. However, previous work has demonstrated that continuous theta burst stimulation (cTBS) [18] can modulate the activity of regions within the larger functional network of the stimulation site [19–25]. Here we adopted this approach by administering cTBS to a lateral prefrontal cortex (LPFC) coordinate individually determined to have maximal resting-state functional magnetic resonance imaging (rs-fMRI) connectivity with the intended OFC target. Based on previous animal inactivation and lesion studies [6–13], we hypothesized that by targeting a region functionally connected to OFC, we would temporarily disrupt activity in the larger OFC network, and thus selectively impair inference-based choices in the devaluation task.

## RESULTS

### Learning of cue-outcome associations during training

We administered cTBS to two groups of healthy subjects (STIM: N=28, cTBS at 80% resting motor threshold [RMT]; SHAM: N=28, cTBS at 5% RMT) in the context of a reinforcer devaluation task (**Fig. 1A**). In an initial training session, hungry subjects learned associations between visual cues and two individually selected food odor rewards (**Fig. 1B-C**). On the next day, preferences for the two food odors predicted by these cues were assessed in a *Baseline* free choice task. Subjects then received cTBS to the individually selected target site (**Fig. 1D**), followed by feeding to satiety on a meal congruent with one of the two food odors (**Fig. 1A**, **Table 1**). The effect of cTBS on choices for these food odors was then measured in a *Probe* session (**Fig. 1B-C**).

**Figure 1.**
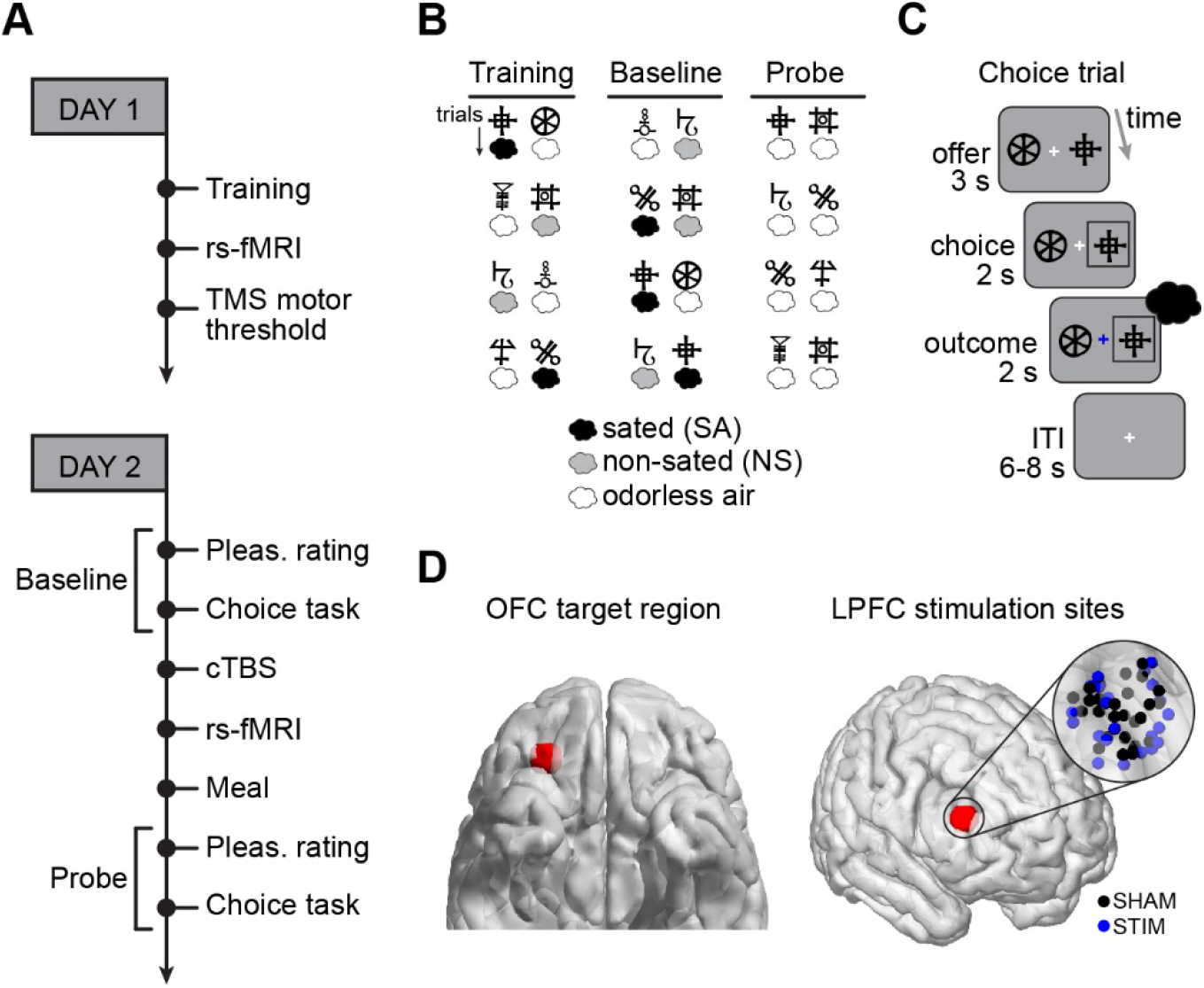
Experimental paradigm and cTBS stimulation sites. **(A)** Day1 and Day2 procedures were conducted on consecutive days. Experimental phases occurring after cTBS on Day2 took place within 1 hour of the end of stimulation (putative duration of the cTBS effect), and there was no difference between STIM and SHAM subjects in the starting time of any phase (*p*’s > 0.44). **(B)** The *Training* session involved choices between 12 unique pairs of visual cues. In 6 pairs, one cue was deterministically paired with the sated odor (SA, black air puff symbol, corresponding to the consumed meal), and the other cue was paired with odorless air (white air puff). In the other 6 pairs, one cue was deterministically paired with the non-sated odor (NS, gray air puff), and the other cue was paired with odorless air. The *Baseline* choice task involved 48 consecutive trials: 24 original pairs, and 24 new pairs in which one cue was associated with the SA odor, and the other cue was associated with the NS odor. The *Probe* choice task involved the same number and type of trials as *Baseline*, but was conducted in extinction, such that odorless air was delivered regardless of the chosen symbol. **(C)** The same trial timing was used for choice trials in the *Training*, *Baseline*, and *Probe* sessions. **(D)** Using the Neurosynth database of rs-fMRI data, we identified a coordinate in central/lateral OFC (*x* = 28, *y* = 38, *z* = −16) that has high functional connectivity (*r* > 0.2) with a region of LPFC that is accessible to TMS (centered on *x* = 48, *y* = 38, *z* = 20). Individual stimulation sites (inset, right) were determined as the coordinate within a 4-voxel radius sphere surrounding the LPFC coordinate (red sphere on right image) that has maximal connectivity with activity in a 4-voxel radius sphere surrounding the OFC coordinate (red sphere on left image).

**Table 1.** Demographic information and feeding behavior results (mean ± SD).

|  | **STIM** | **SHAM** | **ACTL** |
| --- | --- | --- | --- |
| **Number of subjects** | 28 | 28 | 10 |
| Male | 12 | 12 | 4 |
| Female | 18 | 18 | 6 |
| **Age (years)** | 24 ± 3.5 | 24 ± 4.5 | 25.9 ± 4.1 |
| **Body Mass Index (BMI)** | 23.0 ± 3.4 | 24.2 ± 4.8 | 24.2 ± 4.9 |
| **Calories Consumed total**<br>(kcal) | 595.0 ± 178.8 | 553.8 ± 215.3 | 568.3 ± 265.3 |
| Sweet meal | 558.2 ± 143.9 (N=14) | 517.6 ± 229.0<br>(N=14) | 479.64 ± 657 (N=5) |
| Savory meal | 631.8 ± 206.8 (N=14) | 589.9 ± 202.6<br>(N=14) | 657 ± 271.56 (N=5) |
| **Hunger Rating Pre** | 7.5 ± 1.1 | 7.6 ± 1.5 | 8.1 ± 1.5 |
| **Hunger Ratings Post** | 2.1 ± 1.6 | 2.4 ± 1.3 | 3.7 ± 1.9 |
| **Hours Fasted** | 10.7 ± 4.6 | 9.8 ± 4.0 | 8.7 ± 4.5 |

In the *Training* session conducted on Day1, subjects in the STIM and SHAM groups learned the cue-outcome associations equally for both the sated (SA) and non-sated (NS) choice types (3-way ANOVA with time [trial blocks] and condition [SA/NS] as within-subject factors, and group [STIM/SHAM] as a between-subject factor: main effect of time: *F*_3,54_ = 97.2, *p* = 4.97×10^−36^; main effect of group: *F*_1,54_ = 0.72, *p* = 0.40; group × time interaction: *F*_3,162_ = 0.58, *p* = 0.63; group × time × condition interaction: *F*_3,162_ = 1.42, *p* = 0.24; **Fig. 2A**).

**Figure 2.**
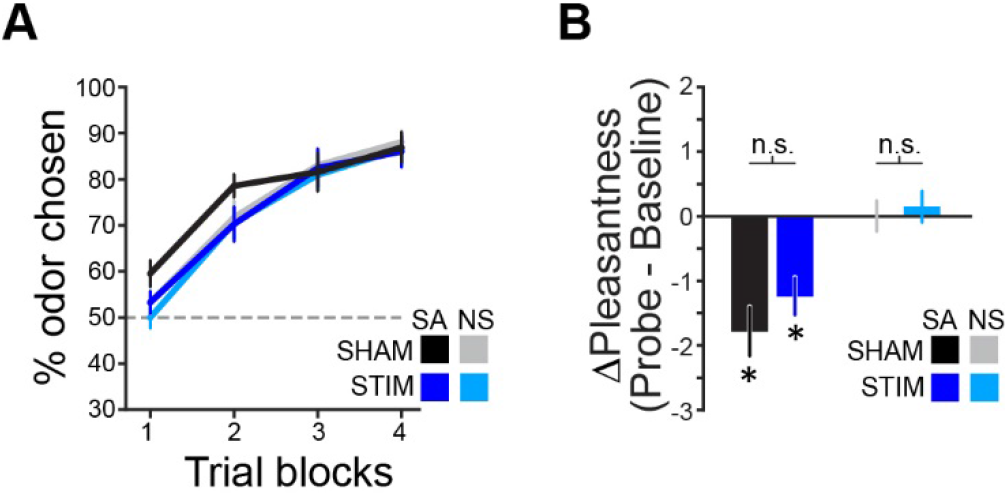
Learning and selective devaluation. **(A)** In the *Training* task, learning was measured as the percentage of trials in which the cue predicting an odor was chosen within each trial block (12 trials per condition per block). Learning was well above 50% chance in the final trial block for both conditions within each group (SHAM: SA *t*_27_ = 11.0, *p* = 1.82×10^−11^, NS *t*_27_ = 15.4, *p* = 7.11×10^−15^; STIM: SA *t*_27_ = 10.4, *p* = 5.87×10^−11^, NS *t*_27_ = 10.6, *p* = 4.30×10^−11^, one-sample *t*-tests), and there was no difference between groups in % odor chosen for either condition (SHAM vs. STIM, SA: *t*_54_ = 0.19, *p* = 0.85; SHAM vs. STIM, NS: *t*_54_ = 0.35, *p* = 0.73, two-sample *t*-tests). Error bars depict within-subject s.e.m. **(B)** There was a significant decrease in pleasantness rating for the SA odor in both the STIM and SHAM groups (asterisks on bar plots, statistics reported in main text), and no change in pleasantness for the NS odor in either group. Error bars depict s.e.m.

### Selective devaluation of food odors

To assess whether consumption of the meal corresponding to one of the two food odors resulted in selective devaluation of that odor, we acquired pleasantness ratings for both odors at the beginning of the *Baseline* and *Probe* phases of the experiment on Day2. There was a significant interaction between condition (SA/NS) and session (*Baseline*/*Probe*) on pleasantness ratings (3-way ANOVA, *F*_1,54_ = 34.6, *p* = 2.60×10^−7^), but no main effect of group (*F*_1,54_ = 2.36, *p* = 0.13) or interaction involving group (group × condition: *F*_1,54_ = 1.10, *p* = 0.30; group × session: *F*_1,54_ = 1.17, *p* = 0.28; group × condition × session: *F*_1,54_ = 0.54, *p* = 0.46; **Fig. 2B**). Follow-up 2-way ANOVAs revealed significant interactions between condition and session in both groups (SHAM: *F*_1,27_ = 22.3, *p* = 6.42×10^−5^; STIM: *F*_1,27_ = 13.0, *p* = 0.0012), which were driven by a decrease in pleasantness for the sated odor (SHAM: *t*_27_ = 4.69, *p* = 7.02×10^−6^; STIM: *t*_27_ = 4.29, *p* = 2.02×10^−4^, paired *t*-tests), and no change in pleasantness for the non-sated odor (SHAM: *t*_27_ = 0.02, *p* = 0.99; STIM: *t*_27_ = 0.60, *p* = 0.55, paired *t*-tests). Thus, consistent with prior work, disruption of OFC activity did not affect the ability to update the value of rewards themselves [11, 12, 26].

### OFC-targeted cTBS disrupts choices for devalued outcomes

We next tested whether targeted OFC stimulation had an effect on subjects’ ability to infer that new value to adapt their choice behavior. In a comparison of choices made in the *Baseline* session to those made in the earliest trials of the *Probe* session, there was an interaction between group and session on the percentage of trials in which the sated odor was chosen (2-way ANOVA: *F*_1,54_ = 8.03, *p* = 0.0064; **Fig. 3A**). This effect was driven by a significant decrease in choices for the sated odor after devaluation in the SHAM group (*t*_27_ = 4.23, *p* = 2.37×10^−4^, paired *t*-test, *Baseline* vs. 1^st^ *Probe* block) and no change in responding in the STIM group (*t*_27_ = 1.34, *p* = 0.19, paired *t*-test, *Baseline* vs. 1^st^ *Probe* block; **Fig. 3B**). Thus while subjects in the SHAM group redirected choices away from cues predicting the devalued odor, subjects in the STIM group failed to show this effect of selective devaluation on choices, and continued to respond at the same rate as in *Baseline*. This group difference was also evident on the very first trial of the *Probe* session (% sated odor chosen, SHAM vs. STIM: *t*_54_ = −2.44, *p* = 0.0176, two-sample *t*-test; **Fig. 3**), further demonstrating that OFC-targeted cTBS impaired the ability to infer the new value of the devalued outcome.

**Figure 3.**
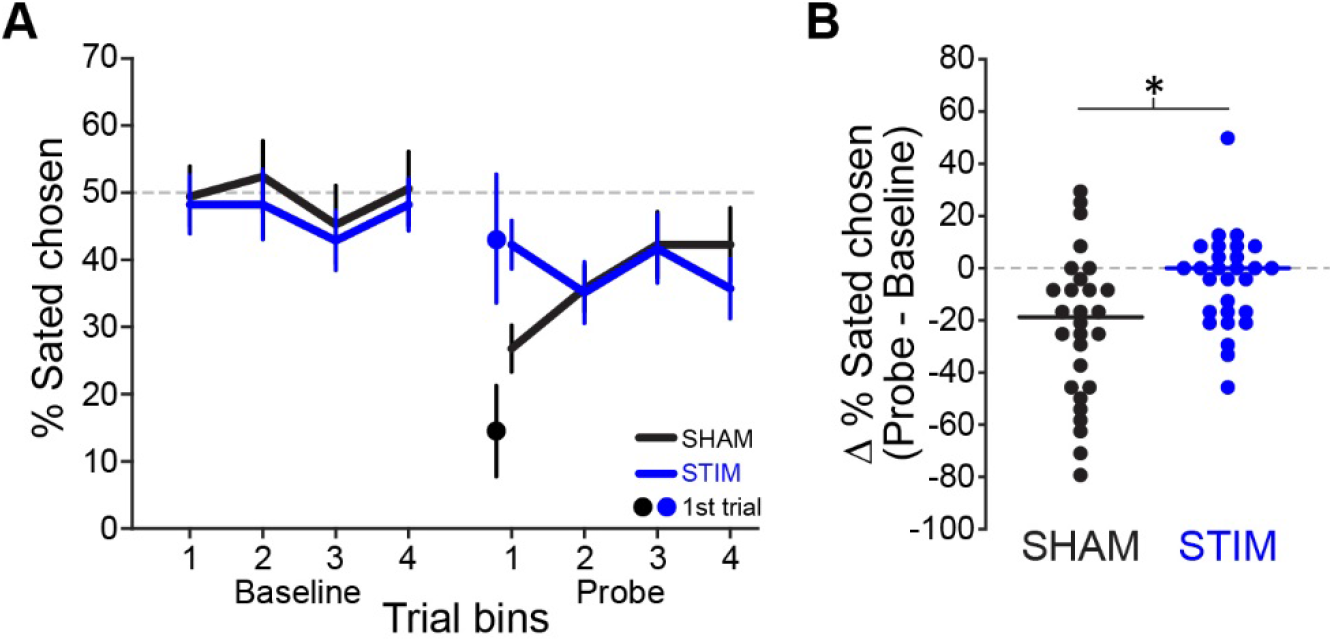
OFC-targeted cTBS impairs inference-based choices. **(A)** The percentage of choice trials in which the sated odor was chosen is plotted for each trial bin (6 trials per bin) in the *Baseline* and *Probe* session. Percent sated odor chosen averaged across *Baseline* trial bins was not different between the groups (SHAM vs. STIM, *t*_54_ = 0.47, *p* = 0.64, two-sample *t*-test) and was not different from 50% in either group (SHAM: *t*_27_ = 0.13, *p* = 0.89; STIM: *t*_27_ = 0.96, *p* = 0.34, one-sample *t*-tests). **(B)** The change in % sated odor chosen from *Baseline* to the first *Probe* trial bin is plotted for individual subjects (each circle = 1 subject). The solid line depicts the median within each group.

### OFC-targeted cTBS reduces global connectedness of OFC

To characterize the effects of OFC-targeted cTBS on OFC network activity, we analyzed rs-fMRI data acquired the day before (Day1) and immediately after (Day2) stimulation. For this, we first calculated a measure of absolute “connectedness” between each voxel’s time series of activity and the rest of the brain, and then computed the change in connectedness from Day1 to Day2 to generate subject-specific difference maps (**STAR Methods**). We then conducted a group-level analysis, comparing these difference maps between the STIM and SHAM group. This analysis revealed a focal effect of stimulation on connectedness in OFC (*x* = 34, *y* = 50, *z* = −8, *p* = 0.00036; **Figure 4A**). *Post hoc* tests confirmed that the significant group effect in OFC was driven by reduced OFC network connectivity in the STIM group (*Z* = 2.30, *p* = 0.021, Wilcoxon signed rank test), whereas no changes were found in the SHAM group (*Z* = 1.34, *p* = 0.18, Wilcoxon signed rank test; **Figure 4B**).

**Figure 4.**
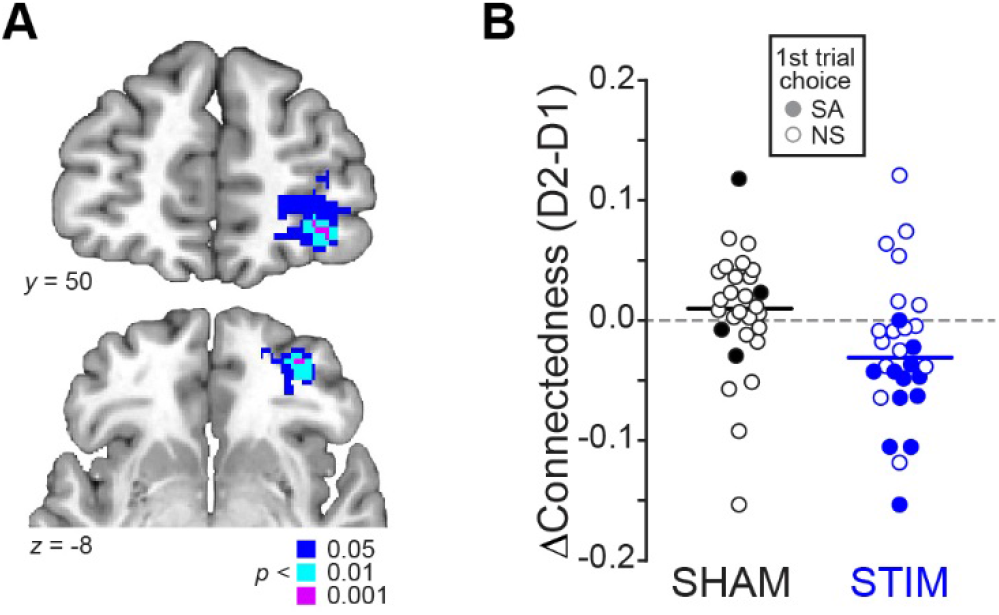
OFC-targeted cTBS disrupts OFC network activity. **(A)** Coronal (top) and axial (bottom) slices show voxels exhibiting a significant interaction between group (STIM/SHAM) and rs-fMRI scanning session (D1/D2) on whole-brain connectedness. Effects are shown at *p* < 0.05 (blue), *p* < 0.01 (cyan), and *p* < 0.001 (magenta), uncorrected for illustration. **(B)** Change in connectedness in each group is shown in individual subjects. Filled circles depict subjects who chose the cue predicting the sated odor in the first trial of the *Probe* session, and empty circles depict subjects who chose the cue predicting the non-sated odor.

We next asked whether the significant change in connectedness in the STIM group was related to the behavioral impairment observed in the choice task. We hypothesized that if behavioral changes were related to changes in OFC connectivity, stronger reductions in OFC connectedness should be accompanied by a higher probability of selecting the cue associated with the devalued outcome in the probe test. In line with this prediction, we found that subjects in the STIM group with a larger reduction in OFC network connectivity (median split) were more likely to choose the cue predicting the devalued odor (*Χ*^*2*^_*1*_ = 9.33, *p* = 0.0023, Chi-square test; **Figure 4B**). There was no comparable relationship between OFC connectivity and choice behavior in the SHAM group (*Χ*^*2*^_*1*_ = 0, *p* = 0.99, Chi-square test). These results provide evidence for a direct relationship between the effect of cTBS on OFC and the effect of cTBS on choice behavior, suggesting that OFC network activity is necessary for outcome-guided behavior. Importantly, these effects were specific to the OFC; there was no effect of cTBS on global connectivity at the individually determined stimulation sites in LPFC (STIM group: *Z* = 1.39, *p* = 0.16, Wilcoxon signed rank test), and no relationship between choice behavior and connectedness at those sites (*Χ*^*2*^_*1*_ = 0.58, *p* = 0.44, Chi-square test).

### OFC-targeted cTBS does not disrupt choices in general

It is possible that the observed effect of cTBS on inference-based choices in the STIM group was due to a more general disruption of behavior. That is, STIM subjects might have been unable to discriminate the cues or to access any value representation, and so may have been responding randomly in the *Probe* session. To rule out this possibility, we analyzed behavior on trials involving choices between cues predicting an odor and odorless air (**Figure 1B**). In a 3-way ANOVA, there was no interaction between group, session, and condition on the percentage of trials in which odor was chosen (*F*_1,54_ = 0.035, *p* = 0.85). Follow-up tests revealed that percentage odor chosen was above chance in the *Baseline* session for both conditions in both groups (SHAM, SA: *t*_27_ = 6.80, *p* = 2.62×10^−7^; SHAM, NS: *t*_27_ = 6.12, *p* = 1.56×10^−6^; STIM, SA: sated: *t*_27_ = 5.56, *p* = 6.80×10^−6^; STIM, NS: *t*_27_ = 7.15, *p* = 1.09×10^−7^, paired *t*-tests) and remained above chance in the first trial block of the *Probe* session (SHAM, SA: *t*_27_ = 2.58, *p* = 0.016; SHAM, NS: *t*_27_ = 6.80, *p* = 2.62×10^−7^; STIM, SA: *t*_27_ = 3.10, *p* = 0.0045; STIM, NS: *t*_27_ = 6.02, *p* = 2.01×10^−6^, paired *t*-tests; **Figure S1**). These data show that subjects in the STIM group were not responding randomly, indicating that cTBS did not disrupt general perceptual or choice-related functions.

### Outcome-guided choices are not affected by unspecific effects of TMS to LPFC

Another possibility is that our results were driven by unspecific effects of cTBS, such as stress or anxiety caused by incidental stimulation of facial muscles and general discomfort associated with cTBS to frontal areas. To rule this out, we repeated the experiment in an independent sample (N=10) using an active control (ACTL) stimulation protocol, designed to induce comparable levels of facial muscle movement and general discomfort, but without inducing changes in underlying neural activity (**STAR Methods**). Subjects in the ACTL group learned the initial cue-outcome associations (% odor chosen in final learning block vs. chance, SA: *t*_9_ = 4.53, *p* = 0.0014; NS: *t*_9_ = 12.5, *p* = 5.32×10^−7^, paired *t*-tests; **Fig. 5A**), and showed selective devaluation of the odor related to the consumed meal (2-way ANOVA, session × condition interaction: *F*_1,9_ = 17.0, *p* =0.0026; driven by a change in pleasantness for the SA odor [*t*_9_ = 4.71, *p* = 0.0011], and no change for the NS odor [*t*_9_ = 0.50, *p* = 0.63]; **Fig. 5B**). 3-way ANOVAs indicate learning and devaluation were comparable to SHAM and STIM subjects (Learning, group × time × condition interaction: *F*_6,189_ = 0.92, *p* = 0.48; Devaluation, group × session × condition interaction: *F*_2,63_ = 1.46, *p* = 0.24).

**Figure 5.**
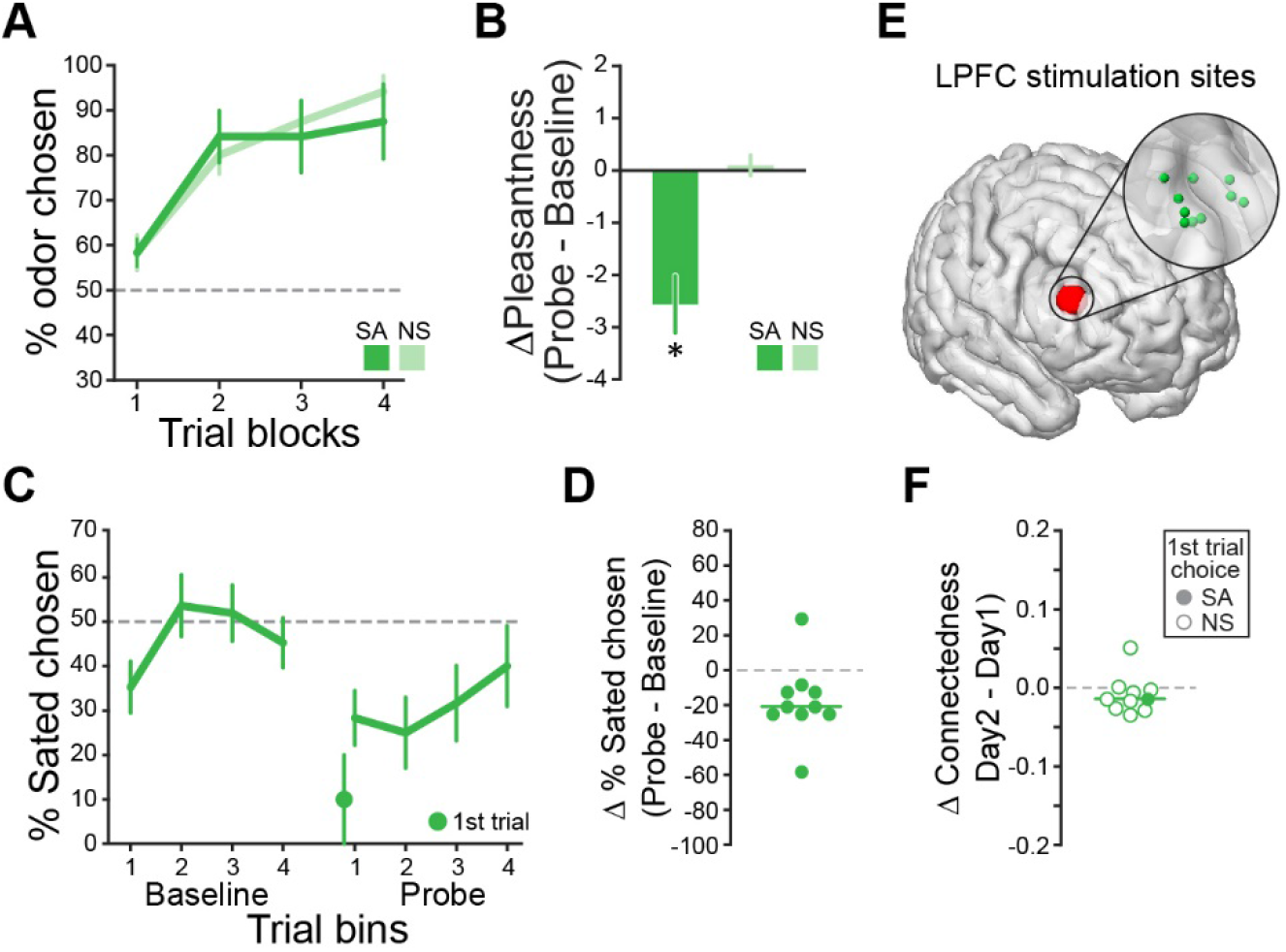
Behavior in an active control stimulation group resembles that of SHAM subjects. **(A)** ACTL subjects showed levels of initial learning in the *Training* session and selective devaluation **(B)** comparable to SHAM and STIM subjects. Error bars depict s.e.m. **(C)** ACTL subjects showed a significant effect of devaluation on choice behavior, such that they chose the SA odor significantly less in the first block of the *Probe* session compared to *Baseline*. Error bars depict s.e.m. **(D)** Effect of devaluation on choices is shown for individual subjects. **(E)** Individual sites for ACTL stimulation in LPFC were determined in the same manner as was done for cTBS stimulation. **(F)** In the OFC region that exhibited a significant change in connectedness after cTBS in the STIM group, there was no change in connectedness in the ACTL group. Each circle represents a subject. Filled circles depict subjects who chose the cue predicting the sated odor in the first trial of the *Probe* session, and empty circles depict subjects who chose the cue predicting the non-sated odor.

Most importantly, ACTL subjects showed a significant effect of devaluation on their choice behavior (% SA chosen, mean *Baseline* vs. first *Probe* trial bin: *t*_9_ = 2.63, *p* = 0.027, paired *t*-test; % SA odor chosen on 1^st^ *Probe* trial vs. chance: *t*_9_ = 4.00, *p* = 0.0031, one-sample *t*-test; **Figure 5C**). This effect was significantly different from the STIM group (*t*_36_ = 1.89, *p* = 0.033, one-tailed, two-sample *t*-test), but similar to the SHAM group (*t*_36_ = 0.48, *p* = 0.64, two-sample *t*-test). Finally, we found that the ACTL stimulation had no effect on connectedness in the same OFC region observed in the STIM group (*Z* = 1.40, *p* = 0.16, Wilcoxon signed rank test; **Fig. 5E-F**), and OFC connectivity was not related to choices in the probe test (*Χ*^*2*^_*1*_ = 1.11, *p* = 0.29, Chi-square test). Together, results from this control experiment suggest that unspecific effects of stimulation are very unlikely to account for the behavioral effects observed with cTBS.

## DISCUSSION

The primary contribution of OFC to decision making has been a matter of long-standing debate [27]. Prominent theories postulate that OFC is necessary for response inhibition [28], representing somatic markers [29], storing stimulus-outcome associations [30], prediction errors [31], credit assignment [32], signaling specific outcome expectations [33], or computing economic value [34]. This diversity of proposals is reflected in the heterogeneity of decision-related signals encoded in this region [35–47], even in individual studies. For instance, a recent electrophysiological recording study in human neurosurgery patients found that a variety of choice and outcome variables, such as value, risk, and regret, were correlated with OFC activity [48].

In the face of such promiscuous neural coding, studies that use experimental lesions or reversible disruption of activity are indispensable for providing a clearer picture of its critical contribution. By administering non-invasive OFC-targeted stimulation in the context of a devaluation task, here we provide evidence for a specific causal role for OFC in outcome-guided behavior in healthy humans, echoing previous work in rats [6–9], non-human primates [10–13, 49], and human patients with lesions encompassing this area [50]. These studies all converge on the finding that OFC is critical for flexibly linking predictive cues to expected rewards and their current value.

Our results are also compatible with previous human imaging [14–16] and animal recording studies using devaluation tasks [51, 52], indicating that OFC activity is specifically modulated in response to cues predicting devalued rewards. Together with the lesion studies cited above, these results suggest that OFC is critical for value-based decision making, but only when the value of specific outcomes has to be inferred [14, 27, 53]. It is possible that value is just one of many potentially relevant features of expected outcomes, including their timing, probability, and sensory properties, that together make up a cognitive map of task space that enables the model-based simulation or inference of future outcomes [54–56]. This theoretical framework can reconcile the multitude of decision-related signals previously found in the OFC.

Because the OFC is not directly accessible to TMS, we applied stimulation to a site in the LPFC that is maximally connected to the intended OFC target. This approach has previously been used to modulate activity in downstream areas connected to the stimulation site, and has been shown to change behavior and functions that depend on these downstream areas [19–25]. However, on its face, it is possible that the behavioral effects observed here are due to activity changes in the LPFC rather than the OFC. We believe this is unlikely for several reasons. First, the connectedness analysis only identified effects of cTBS in the OFC but not in the LPFC. Second, the behavioral effects of cTBS were directly related to effects of cTBS on OFC network connectivity but not on LPFC connectivity. Third, we did not find effects in our ACTL group who received active stimulation to the same individually selected LPFC area, albeit at a different stimulation frequency, which is not expected to cause effects in downstream targets. Finally, while multiple animal studies across different species have shown that OFC is necessary for responding in the reinforcer devaluation task [6–13], we are not aware of comparable positive findings in the LPFC. Taken together, although we cannot rule out the possibility that effects of cTBS on LPFC activity contributed to the behavioral impairment, we are confident that cTBS-induced modulation of OFC network connectivity was a significant factor.

It is important to note that our results provide evidence for the feasibility of targeting human OFC with non-invasive stimulation, thereby highlighting the potential of this technique to study the role of OFC in health and to modulate its function in disease. Disruption in OFC function is implicated in a variety of neurological and neuropsychiatric conditions, including depression [57, 58], obsessive compulsive disorder [59, 60], and substance abuse [61–63], and microstimulation of these networks has been shown to restore drug-induced behavioral deficits in animal models of addiction [64, 65]. Our results thus provide the basis for the development of novel stimulation protocols targeting OFC networks in humans to treat such disorders [17].

## Acknowledgments

The authors thank International Flavor and Fragrances (R.S. Santos) and Kerry (J.L. Buckley) for providing food odorants. This work was support by National Institute on Deafness and other Communication Disorders Abuse (NIDCD) grant R01DC015426 (to T.K.) and the Intramural Research Program at the National Institute on Drug Abuse.

## Author Contributions

J.D.H., and T.K. conceived the study and designed the experiments with input from J.L.V. and G.S.. J.D.H, D.E.S., and R.R. collected the data. J.D.H analyzed the data. T.K. supervised the project. J.D.H., J.L.V., G.S., and T.K. wrote and revised the manuscript. The opinions expressed in this article are the authors’ own and do not reflect the view of the NIH/DHHS.

## METHODS

### Subjects

A total of 89 subjects participated in the initial screening session (see <u>*Experimental design*</u> below). Of these, 56 subjects further participated in the main experiment and were randomly assigned to either the SHAM (*n* = 28, 16 female) or STIM (*n* = 28, 16 female). After the main experiment was conducted, an independent group of these subjects participated in the active control experiment (ACTL, *n* = 10, 6 female). For demographic and other behavioral information by group, see **Table 1**. All subjects provided written consent to participate, reported no neurological or psychiatric disorders, no history of seizures, and were not currently taking psychotropic drugs. Eligibility for transcranial magnetic stimulation (TMS) was determined based on standardized safety guidelines [66]. Subjects were compensated with $20 per h for behavioral testing, and $40 per h for MRI scanning and TMS. The study was approved by the Northwestern University Institutional Review Board.

### Odor stimuli and presentation

Eight food odors, including four sweet (pineapple cake, caramel, strawberry, gingerbread) and four savory (potato chips, pot roast, pizza, garlic), were provided by International Flavors and Fragrances (New York, NY) and Kerry (Melrose Park, IL). For all tasks, odors were delivered to participants’ noses using a custom-built computer-controlled olfactometer capable of redirecting medical grade air with precise timing at a constant flow rate of 3.2 L/min through the headspace of amber bottles containing liquid solutions of the food odors. The olfactometer is equipped with two independent mass flow controllers (Alicat, Tucson, AZ), allowing for dilution of odorants with odorless air. There was a constant stream of odorless air delivered throughout the experiment, and odorized air was mixed into this airstream at specific time points, with no change in the overall flow rate. Thus, odor presentation did not involve a change in somatosensory stimulation.

### Food items

For the meal phase of the main experiment, food items with a dominant flavor note corresponding to one of the two odors selected for each participant were provided for consumption. These food items were as follows: pineapple cake odor: pineapple flavored cakes; caramel odor: caramel sauce on biscuits; strawberry odor: strawberry wafers; gingerbread odor: gingersnap cookies; potato chip odor: potato chips; pot roast odor: pot roast; pizza odor: cheese pizza; garlic odor: garlic bread. All food items were procured from Whole Foods, Trader Joe’s, H Mart, or Jewel Osco.

### Experimental design

The experiment consisted of an initial screening session conducted in a behavioral testing room adjacent to the main lab space, followed by two consecutive days of experimental sessions (*Day 1* and *Day 2*) conducted at a later date in rooms available at the MRI scanning facility. The *Day 1* session of the main experiment was conducted on average 18.4 days (± 1.77 days, s.e.m.) after the screening session. For all sessions, subjects were instructed to arrive in a hungry state, having fasted for at least 4-6 h prior to testing. Odor pleasantness ratings were made on a visual analog scale using a scroll wheel and mouse button press. Pleasantness rating anchors were “most liked sensation imaginable” and “most disliked sensation imaginable”.

#### Screening session

Subjects first rated the pleasantness of the 8 food odors. Based on visual inspection of these ratings by the experimenter, one sweet odor and one savory odor were selected such that they were both rated as pleasant (i.e., above the “neutral” line on pleasantness scale), and matched as closely as possible in their rating. These 2 selected odors were then used as unconditioned stimuli for that individual subject for the remainder of the experiment. If these criteria were not met (e.g., if none of the 4 savory odors were rated above neutral in pleasantness), the subject was excluded from further participation in the experiment. Combined with subjects who “passed” the screening but were not available for scheduling of the main experiment at a later date, a total of 23 of the 89 subjects who participated in the screening session did not further participate in the *Day 1* and *Day 2* sessions described below.

##### Day 1

In a behavioral testing room adjacent to the MRI scanner, subjects first completed a training choice task to learn associations between abstract visual symbols and odor outcomes. This task consisted of 12 unique pairs of visual cues, randomly chosen for each subject independently. Within each pair, one cue was associated with an odor outcome, and one was associated with odorless air. Six pairs were associated with the sweet odor, and the other 6 were associated with the savory odor. On each trial of the task, the two cues in a given pair were presented on the screen simultaneously to the left and right of a white center crosshair. Subjects had 3 s to make a left or right mouse button click to choose the corresponding cue. The chosen cue was then highlighted, and after a 2 s delay the center crosshair turned blue, indicating that the outcome associated with the chosen symbol was present and they should make a sniff. The training task consisted of 4 blocks of 24 trials each, with each pair presented twice per block (left/right position of cue pairs counterbalanced). Prior to the training task, subjects were instructed to learn which of the two cues in each pair led to an odor outcome, and to choose those symbols.

After the training task, we acquired a structural T1-weighted MRI scan to aid in anatomical guidance of TMS. We also acquired an 8.5-minute baseline resting state fMRI (rs-fMRI) scan, which was used to identify the specific coordinate at which to apply cTBS on the following day (see *TMS target coordinate selection* below). In a room dedicated for TMS adjacent to the MRI scanner, we then determined resting motor threshold (RMT) (see *Transcranial Magnetic Stimulation* below).

##### Day 2

Subjects first completed a *Baseline* behavioral session consisting of pleasantness ratings of the food odors and a choice task. The choice task consisted of 48 consecutive choice trials using the same trial timing described above for the training task. Twenty four trials in this task were the original odor/odorless pairs learned on the previous day, and the remaining 24 trials were new pairs consisting of one cue associated with the sweet odor and one cue associated with the savory odor. The trial order was pseudorandomized such that 12 original (6 sweet/odorless, 6 savory/odorless) and 12 new (sweet/savory) trials were presented in random order within each half of the task. Subjects were instructed that this was a free choice task, and they should choose whichever of the two symbols they wanted based on the odor outcome they expected to receive.

After the *Baseline* session, subjects received cTBS (STIM group: 80% RMT; SHAM group: 5% RMT). Immediately after the stimulation, we acquired another 8.5-minute rs-fMRI scan. In a separate testing room adjacent to the scanner, subjects were then given a meal with a dominant flavor note corresponding to one of the two food odors used in the experiment (pseudorandomized). For this meal phase, subjects were instructed to eat as much as they wanted within a 15-minute time period. Hunger ratings between 0 and 10 (0 = “not at all hungry”, 10 = “extremely hungry”) were acquired before and after the meal.

After the meal, subjects completed a *Probe* behavioral testing session consisting first of odor pleasantness ratings, and then 48 choice trials in extinction (i.e., odorless air was delivered regardless of the choice). The same pseudo-randomization of choice trials was used as described above for the *Baseline task*, except that the first 3 trials were always sweet/savory pairs.

### Transcranial Magnetic Stimulation

We used a MagPro X100 stimulator connected to a MagPro Cool-B65 butterfly coil (MagVenture A/S, Farum, Denmark) to deliver TMS guided anatomically by the individual T1-weighted anatomical scans acquired on *Day 1*. Stimulation was administered in a room designated for TMS adjacent to the MRI scanner. For determination of RMT, we delivered single pulses starting at 50% of maximum stimulator output over left motor cortex, and adjusted stimulation strength as necessary to locate a site that evoked isolated movements of the right thumb. At this location, RMT was determined as the minimum percentage of stimulator output necessary to evoke 5 visible thumb movements in 10 stimulations.

The cTBS protocol on *Day 2* lasted 40 s and consisted of 600 total pulses delivered at either 80% RMT (STIM group) or 5% RMT (SHAM group). Each burst in this sequence included 3 pulses delivered at 50 Hz, and bursts occurred every 200 ms (5 Hz) [18]. The active control (ACTL) stimulation lasted 7.5 m and consisted of a total of 600 pulses delivered at 20 Hz in 2 s trains, with 28 s of no stimulation between pulse trains. ACTL stimulation was delivered at approximately 50% RMT, which was the limit of tolerability as determined by 2 s test trains delivered to the stimulation site prior to administration of the full 7.5 m stimulation sequence. Because of the length of the pulse trains in the ACTL sequence, these pulses caused comparatively more facial muscle movement and discomfort than the cTBS sequence, and therefore resulted in the decreased level of stimulation. However, even at approximately 50% RMT, the ACTL sequence still caused levels of facial muscle movement comparable to cTBS at 80% RMT. This stimulation is thus an appropriate control for the possible effects of stress or discomfort on subsequent task performance.

Both cTBS and ACTL stimulation were applied at the coordinate in lateral prefrontal cortex determined individually to have maximal functional connectivity with the orbitofrontal cortex seed coordinate (see <u>*TMS target coordinate selection*</u> below). All subjects were informed that stimulation might cause muscle twitches in the forehead, eye area, and jaw. To demonstrate this potential movement and test for tolerability of stimulation at this location, we administered two test pulses. One subject originally designated to be in the STIM group did not tolerate the test pulses, and was thus administered sham stimulation and moved to the SHAM group (all results reported here remain significant even if this subject is excluded). Immediately after the last pulse the time was noted, and starting times of subsequent experimental phases were calculated in reference to this time. All subsequent phases took place within 1 hour of the end of stimulation.

### MRI data acquisition

MRI data were acquired on a Siemens 3T PRISMA system equipped with a 64-channel head-neck coil. For resting state fMRI, echo-planar imaging (EPI) volumes were acquired with a parallel imaging sequence with the following parameters: repetition time, 2 s; echo time, 22 ms; flip angle, 80°; multi-band acceleration factor, 2; slice thickness, 2 mm, no gap; number of slices, 58; interleaved slice acquisition order; matrix size, 104 × 96 voxels; field of view 208 mm × 192 mm. The functional scanning window was tilted ~30° from axial to minimize susceptibility artifacts in OFC [67, 68]. Each fMRI session (*Day 1* and *Day 2*) consisted of 250 EPI volumes covering all but the most dorsal portion of the parietal lobes. On *Day 1*, a 1 mm isotropic T1-weighted structural scan was also acquired for navigation of stimulation and to aid in spatial normalization.

### fMRI data preprocessing

Image preprocessing was performed using SPM12 software (www.fil.ion.ucl.ac.uk/spm/). To correct for head motion during scanning, images acquired in the *Day 1* and *Day 2* rs-fMRI session were aligned to the first acquired image in each session. The mean realigned images for each session were then co-registered to the T1 scan, and the resulting registration parameters were applied to the realigned EPI’s. The T1 image was normalized to Montreal Neurological Institute (MNI) space using the 6-tissue probability map provided by SPM12 to generate forward and inverse deformation fields. For TMS target coordinate selection, the co-registered EPI’s corresponding to the *Day 1* session were smoothed with a 6 × 6 × 6 mm Gaussian kernel. For the group-level connectedness analysis described below, the realigned and co-registered *Day 1* and *Day 2* scans were normalized to MNI space using the forward deformation fields generated by normalization of the T1 image. The normalized *Day 1* and *Day 2* scans were smoothed using a 6 × 6 × 6 mm Gaussian kernel.

### TMS target coordinate selection

We used the Neurosynth (www.neurosynth.org) database of rs-fMRI scans to select a coordinate that is both in the vicinity of the central/lateral portion of OFC that has been previously implicated in outcome-guided behavior [10–12, 14], and has high functional connectivity to a surface location that is directly accessible to TMS. This resulted in identification of a coordinate in central/lateral OFC (x=28, y=38, z=−16) that is connected with a coordinate in lateral prefrontal cortex (x=48, y=38, z=20) with a correlation of *r* = 0.26.

For determination of individual stimulation coordinates in LPFC, we first generated spherical masks of 8-mm radius around these two coordinates in MNI space, both inclusively masked by the gray matter tissue probability map provided by SPM12 (thresholded at > 0.1). These masks were un-normalized to each subject’s native space using the inverse deformation field generated by the normalization of the T1 scans. We then specified a general linear model for each subject with the mean *Day 1* rs-fMRI activity in the un-normalized OFC sphere as the regressor of interest (i.e., the seed region), and realignment parameters as regressors-of-no-interest. The stimulation coordinate was calculated as the voxel in the un-normalized LPFC mask that had the highest beta value (i.e., highest functional connectivity with the OFC seed region) estimated from this GLM.

### Global connectedness analysis

For each subject and scanning session (i.e. *Day 1* and *Day 2*), we computed voxel-wise maps of “global connectedness”, reflecting the average connectivity between a given voxel’s time course of rs-fMRI activity and all other gray matter voxels. This was done by first extracting the time course of activity for each voxel in the gray matter tissue probability map mask (threshold at > 0.1). These time courses were then adjusted for head motion by regressing out nuisance parameters, which included: the 6 realignment parameters (3 translations, 3 rotations) calculated for each volume during motion correction; the derivative, square, and the square of the derivative of each realignment parameter; the absolute signal difference between even and odd slices in each volume and the variance across slices in each volume (to account for fMRI signal fluctuation caused by within-scan head motion; additional regressors as needed to model out individual volumes in which particularly strong head motion occurred; the mean global signal in all white matter voxels specified by exclusively masking the white matter tissue probability map with the gray matter tissue probability map. The adjusted time series were then z-scored across scans. We then calculated the absolute Pearson correlation (Fisher’s Z transformed) between each voxel’s time series and every other voxel, resulting in a voxel-by-voxel connectivity matrix. We then averaged across the rows of this matrix, resulting in a measure of global connectedness for each voxel. These whole-brain maps of global connectedness were then compared between days (*Day 2* – *Day 1*) for each subject. Voxels with negative difference values indicate locations in which global connectivity decreased from the *Day 1* baseline scan to the *Day 2* scan acquired immediately after stimulation. In contrast, values close to zero indicate no change in global connectivity. To confirm that cTBS decreased global connectivity of the OFC, we compared these difference maps between groups (median SHAM > STIM) using a permutation test with 100,000 random group assignments.

### Statistics

For testing effects across groups we used mixed-effects ANOVA’s with group as a between-subjects factor and condition, testing session, and trial bins as within-subjects factors. For post hoc testing of effects within groups we used either repeated measures ANOVA or paired *t*-tests. Significance threshold was set to p=0.05, two-tailed, unless otherwise noted.

## Supplementary Figure

**Supplementary Figure 1.**
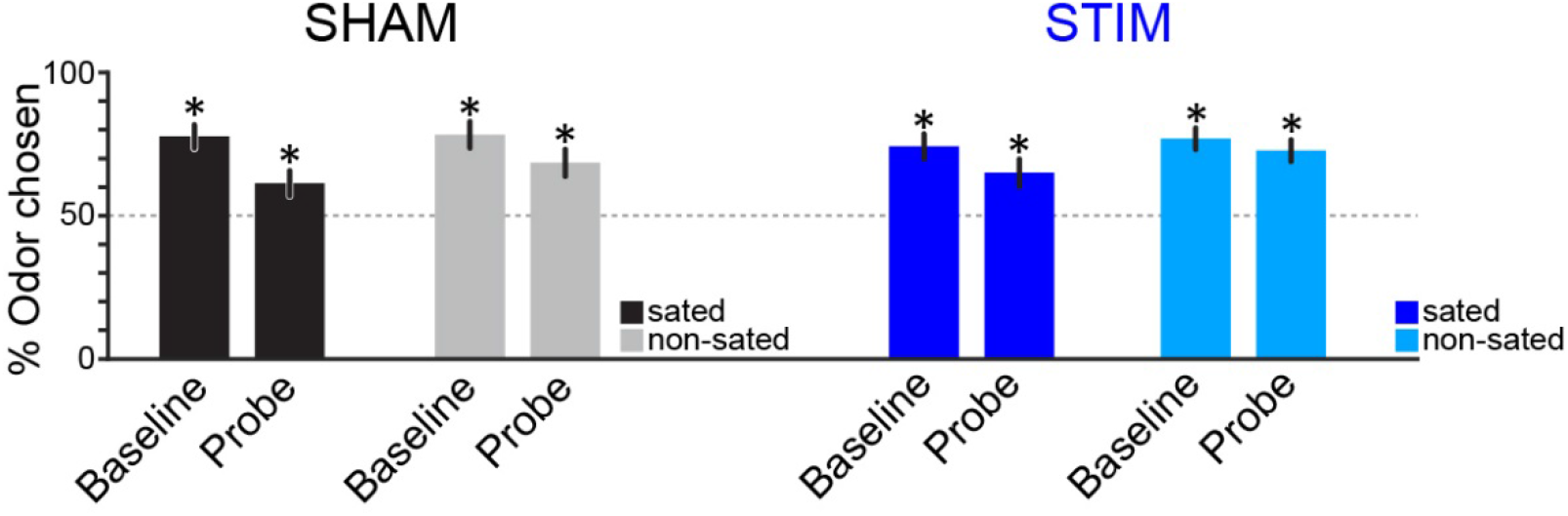
OFC-targeted cTBS does not disrupt choices in general. In the SHAM group, percent odor chosen was above chance in the *Baseline* session for both the sated (*t*_27_ = 6.80, *p* = 2.62×10^−7^) and non-sated (*t*_27_ = 6.12, *p* = 1.56×10^−6^) conditions, and remained above chance in the first trial bin of the *Probe* session (sated: *t*_27_ = 2.58, *p* = 0.016; non-sated: *t*_27_ = 6.80, *p* = 2.62×10^−7^). The same was true in the STIM group, such that percent odor chosen was above chance for both conditions in *Baseline* (sated: *t*_27_ = 5.56, *p* = 6.80×10^−6^; non-sated: *t*_27_ = 7.15, *p* = 1.09×10^−7^) and *Probe* (sated: *t*_27_ = 3.10, *p* = 0.0045; non-sated: *t*_27_ = 6.02, *p* = 2.01×10^−6^) sessions. Subjects were thus not responding randomly, and preferred both sated and non-sated odors over odorless air even after satiety. Error bars depict s.e.m.

